# Allosteric effect of nanobody binding on ligand-specific active states of the β2-Adrenergic Receptor

**DOI:** 10.1101/2021.07.10.451885

**Authors:** Yue Chen, Oliver Fleetwood, Sergio Pérez-Conesa, Lucie Delemotte

## Abstract

Nanobody binding stabilizes the active state of G-protein-coupled receptor (GPCR) and modulates its affinity for bound ligands. However, the atomic level basis for this allosteric regulation remains elusive. Here, we investigate the conformational changes induced by the binding of a nanobody (Nb80) on the active-like β2 adrenergic receptor (β2AR) via enhanced sampling molecular dynamics simulations. Dimensionality reduction analysis shows that Nb80 stabilizes a highly active state of the β2AR with a ~*1*4 Å outward movement of transmembrane helix 6 and close proximity of transmembrane (TM) helices 5 and 7. This is further supported by the residues located at hotspots located on TMs 5, 6 and 7, as shown by supervised machine learning methods. Besides, ligand-specific subtle differences in the conformations assumed by intercellular loop 2 and extracellular loop 2 are captured from the trajectories of various ligand-bound receptors in the presence of Nb80. Dynamic network analysis further reveals that Nb80 binding can enhance the communication between the binding sites of Nb80 and of the ligand. We identify unique allosteric signal transmission mechanisms between the Nb80-binding site and the extracellular domains in presence of full agonist and G-protein biased partial agonist, respectively. Altogether, our results provide insights into the effect of intracellular binding partners on the GPCR activation mechanism, which could be useful for structure-based drug discovery.

**TOC:** Graphical Table of Contents

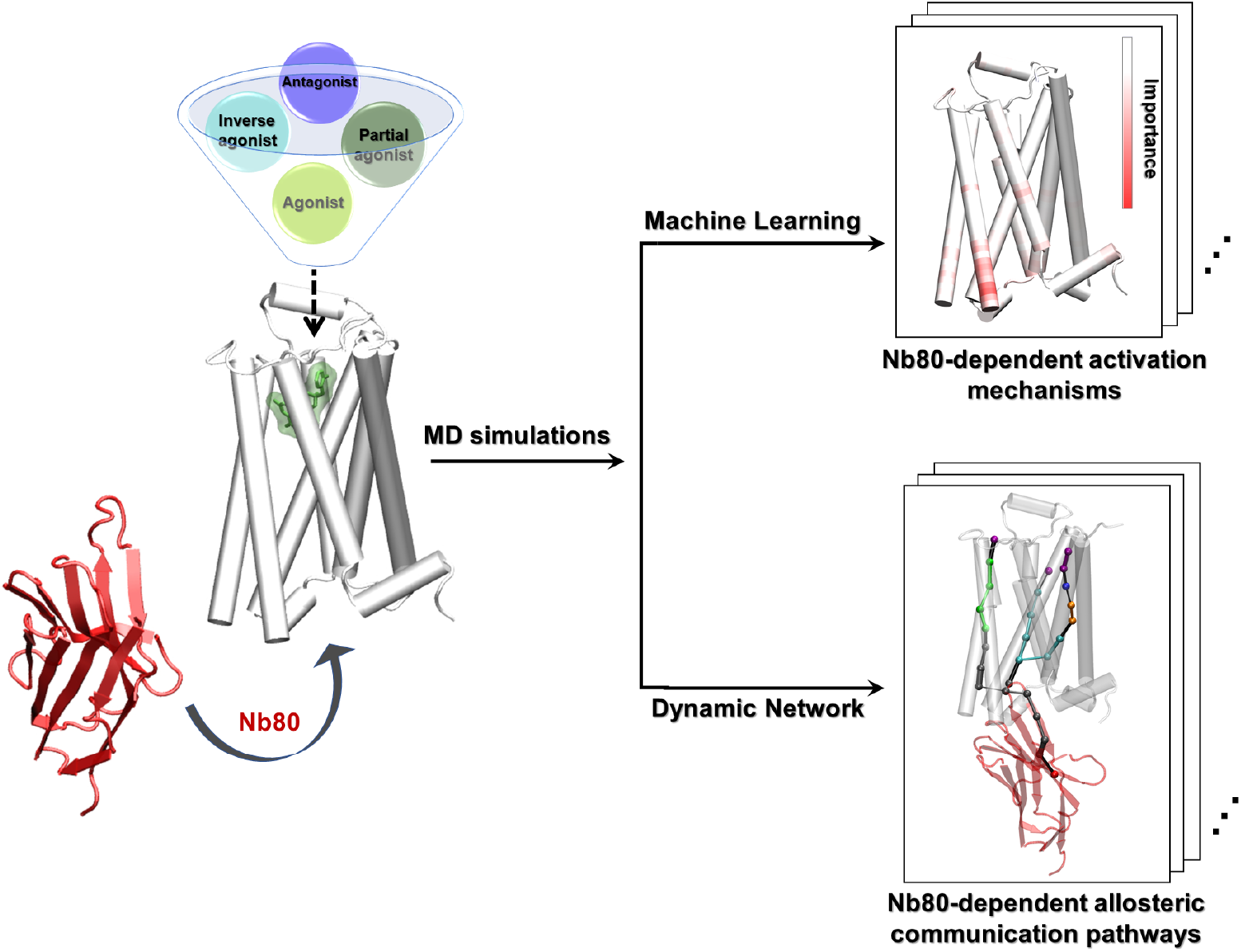

## Introduction

The G-protein-coupled receptor (GPCR) superfamily is the largest and most distinct group of membrane receptors in eukaryotes, comprising over 800 diverse human cell-surface receptors.^1^ They mediate signaling of a variety of extracellular stimuli, including photons, odorants, hormones, peptides and proteins, and regulate many physiological processes. Not surprisingly, GPCRs are important targets for the binding of drugs, which account for ~34% of all US Food and Drug Administration (FDA)-approved medicines, highlighting the indispensable role of GPCRs in health and diseas.^2,3^ All GPCRs share a common seven-transmembrane (7TM) domain helices architecture. They show differences in their extracellular domains, where extracellular ligands bind, and in their intracellular domains, where signaling transducers, such as G proteins and β-arrestins, bind.^4,5^ Upon activation by extracellular ligands, the receptor undergoes certain conformational changes and engages intracellular transducers, which modulates different downstream signaling pathways.

GPCR activation is an allosteric process, involving transducing a signal initiated by various external stimuli into cellular response and downstream regulation of various aspects of human physiology. Therefore, understanding the mechanism underlying the allosteric signaling of GPCRs is of importance for drug discovery and pharmacology research. Several highly conserved residues on the pathway connecting the ligand-binding and the G-protein-binding pockets have been identified by experimental and computational studies.^6,7^ These are organized in microscopic clusters, often referred to as microswitches. Their dynamics play an important role in GPCR activation. For example, the outward movement of transmembrane helix 6 (TM6), located at the cytoplasmic domain of the receptor, is a hallmark of GPCR activation. Some evolutionarily conserved sequence motifs are also identified as microswitches, such as N^7.49^P^7.50^xxY^7.53^ (superscripts referring to Ballesteros-Wein-stein numbering^8^), D^3.49^R^3.50^Y^3.51^, P^5.50^I^3.40^F^6.44^, C^6.47^W^6.48^xP^6.50^, and the sodium-binding pocket at D^2.50^, distributed in the TM domains.^6,7^ In addition, great efforts have also been made to characterize the mechanism underlying the G protein activation,^9,10^ biased signaling,^11,12^ and allosteric modulation^13,14^ via diverse approaches.^15–18^ Several studies, using in particular various spectroscopic techniques, have pointed out that a simple two-state model involving a single inactive and a single active state is an oversimplification and that the activation mechanism instead involves multiple inactive, intermediate and active receptor states.^19,20^ Many studies have focused on the ligand-dependent conformational changes implicated in the enhancement of binding affinity of intracellular transducers.^21,22^ However, the structural basis underlying transducer-induced allosteric communications and how they are related to ligand efficacy is not well understood.

Previously, Fleetwood et al.^23^ focused on analyzing conformational ensembles of the β2 adrenergic receptor (β2AR) modeled in the absence of intracellular binding partner, and revealed that ligands with varying efficacy profiles could stabilize different active-like states of the receptor using enhanced sampling molecular dynamics (MD) simulations coupled with data-driven methods. Here, we build on this work and investigate the structural changes induced by G protein-mimicking Nanobody80 (Nb80)^24^ to the β2AR bound with six ligands with different efficacies (**Figure 1**). This nanobody has been designed, and used, to stabilize the receptor in an active state: the region that comes in interaction with the receptor indeed mimics the interactions made with a G-protein. We first sampled the different active-like ensembles of unliganded and ligand-bound β2AR in the absence and presence of Nb80 using MD simulations. Using dimensionality reduction analysis, we found that Nb80 binding stabilizes a highly active-like state with a 12-14 Å outward movement of TM6 independent of ligand binding. More specifically, BI167107(full agonist)^24^ and salmeterol (G-protein biased partial agonist)^25^ generate different subtle conformational distributions, when compared to the other ligands. In addition to the intracellular end TM6 microswitch, specific residues in TM3, TM5 and TM7 are identified as important features for distinguishing Nb80-bound and N80-free states. In the presence of Nb80, ligand-specific conformational differences mainly show up in the ECL and ICL domains. Furthermore, dynamic network analysis reveals that communication across the receptor is greatly strengthened when binding to Nb80. Interestingly, BI167107- and salmeterol-specific optimal signal transmission pathways from the Nb80-binding site to the ligand-binding site primarily involve TM1 and TM2, and TM5 respectively. Taken together, our findings provide a structural basis for the enhancement of ligand affinity and ligand-specific effects on the receptor activation, controlled by intracellular binding partners, and can potentially be useful in structure-based drug discovery targeting GPCRs.

**Figure 1.**
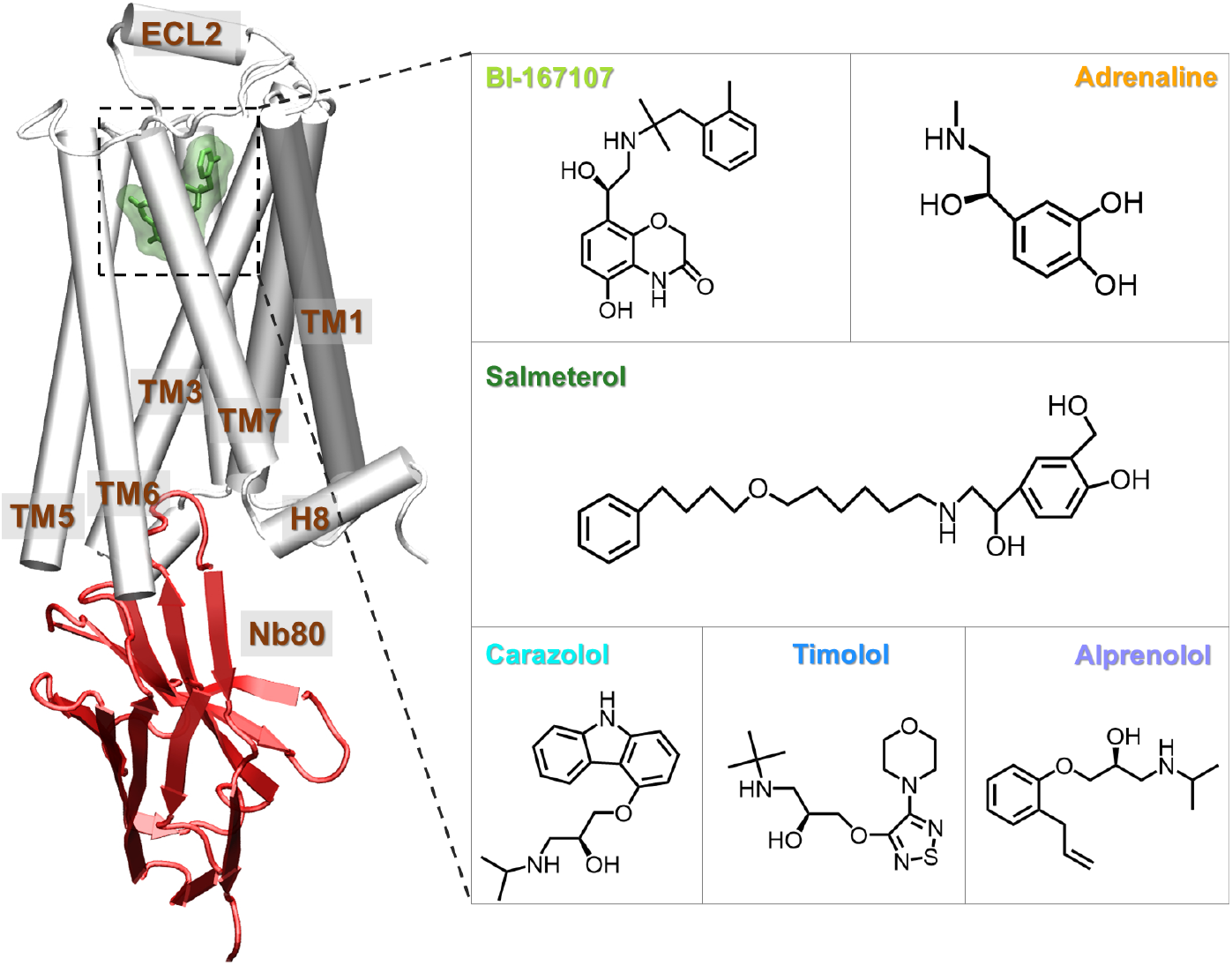
Structure of the β2AR-Nb80: A molecular dynamics snapshot of BI167107-bound β2AR with Nb80 in the active-like state (simulation starting structure: PDB 3P0G^24^), and ligands examined in this study: agonists BI-167107 and adrenaline; biased partial agonist salmeterol; inverse agonist carazolol, and antagonists timolol and alprenolol. The receptor is represented as white cartoon, Nb80 as red ribbons, the bound ligand as green sticks and transparent surface.

## Methods

### Molecular Simulations System Setup

We based all β2AR simulation systems on the active-state structure 3P0G,^24^ which is bound to BI-167107 and Nb80. The nanobody-bound systems had the same system configuration and followed the same equilibration protocol as the previously published nanobody-free simulations.^26,27^ All simulations were initiated with CHARMM-GUI^28^ and used the CHARMM36m force field.^29^ To account for missing residues and mutations present in the experimental structure, we reversed the N187E mutation and capped chain termini with acetyl and methylamide. The ligands included in this study are all resolved in the different β2AR PDB structures 2RH1,^30^ 3NYA,^31^ 3D4S,^32^ 6MXT^33^ and 4LDO.^34^ Due to their localization in hydrophobic environments, E1223.41 and the ligands’ amine groups were protonated, while H172^4.64^ and H178^4.70^ were protonated at their epsilon positions. With a complete model of the β2AR and the Nb80, the protein complex was embedded in a homogeneous POPC^35^ lipid bilayer, and surrounded by a solution consisting of TIP3P water molecules^36^ with a 0.15M concentration of sodium and chloride ions. For the nanobody-bound receptor, we inserted 190 membrane molecules and 120 water molecules per lipid. In the smaller nanobody-free systems, we used 180 membrane molecules and 79 water molecules per lipid. We performed the MD simulations with GROMACS 2018.6.^37^ Input files and simulation trajectories are available publicly online (https://osf.io/b5rav/).^26,27^

### Single state sampling simulations

The conformational ensembles were sampled using kinetically trapped active-like state sampling, or *single state sampling,* a recently published enhanced sampling technique.^27^ In this relatively simple framework, 24 simulation replicas, each of 7.5 ns length, were launched from the starting structure (**Table 1**). Their center point, *c*, was computed in a high-dimensional space spanned by a set of Collective Variables (CVs), which have previously been shown to well characterize the β2AR’s activation mechanism.^27^ For every replica, *i*, we computed its distance to the center, *x_i_*, and the average replica distance to the center, *d*. A weight, 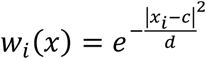, was assigned to every replica. For the next iteration of the method, the replicas were extended, with the number of copies proportional to 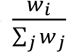 keeping the total replica count at 24. By performing these steps iteratively, the replicas eventually diffused around a well-equilibrated state. In other words, for every ligand-receptor complex, we obtained an ensemble of structures sampled from the closest kinetically stable state accessible from the active starting structure. In line with convergence analysis performed on the original dataset of nanobody-free simulations,^27^ we ran the sampling method for 8 iterations, at which point the center points did not drift between iterations. Finally, the trajectories of the last iteration were further analyzed, as described in the next section.

**Table 1.** Total simulation time per system.

| System | Ligand | Number of iterations | Simulation time (μs) |
| --- | --- | --- | --- |
| apo-β2AR | | 8 | 1.44 |
| BI167107-β2AR | BI167107 | 8 | 1.44 |
| Adrenaline-β2AR | Adrenaline | 8 | 1.44 |
| Salmeterol-β2AR | Salmeterol | 8 | 1.44 |
| Carazolol-β2AR | Carazolol | 8 | 1.44 |
| Timolol-β2AR | Timolol | 8 | 1.44 |
| Alprenolol-β2AR | Alprenolol | 8 | 1.44 |
| apo-β2AR-Nb80 | | 8 | 1.44 |
| BI167107-β2AR-Nb80 | BI167107 | 8 | 1.44 |
| Adrenaline-β2AR-Nb80 | Adrenaline | 8 | 1.44 |
| Salmeterol-β2AR-Nb80 | Salmeterol | 8 | 1.44 |
| Carazolol-β2AR-Nb80 | Carazolol | 8 | 1.44 |
| Timolol-β2AR-Nb80 | Timolol | 8 | 1.44 |
| Alprenolol-β2AR-Nb80 | Alprenolol | 8 | 1.44 |

### Dimensionality reduction methods

We derived the active-like conformational ensemble from the last iteration of the swarms. To project the ensemble onto a lower-dimensional manifold and more easily analyze the active-like state trajectories, we used two different dimensionality reduction methods, principal component analysis (PCA)^38^ and multidimensional scaling (MDS).^39^

PCA is one of the most used techniques for dimensionality reduction. It projects the data on principal components in the linear regime by computing the eigenvectors of the data covariance matrix. The first principal component represents the dimension of the most variance of the data, and the subsequent ones account for decreasing amounts of variance in their respective dimensions. MDS is a non-linear method that includes various multivariate data analysis techniques. It is developed to construct a set of low-dimensional embedding patterns which best preserve pairwise Euclidean distances in the original high-dimensional space.

In this work, we use the PCA and MDS modules of the python package Scikit-learn.^40^ The inverse closest-heavy atom distances were used as input features and the simulation snapshots were then projected onto the first four dimensional feature spaces.

### Supervised feature extraction and learning

We used the supervised feature extraction module implemented in Demystifying,^23^ which aims to identify molecular features that are important for a specific biological question. An artificial feed-forward neural network, a multilayer perceptron (MLP) classifier,^41,42^ was first trained to find the important residues for discriminating β2AR systems in the absence and presence of Nb80. Another MLP was subsequently trained to distinguish all the Nb80-bound systems bound to different ligands. Inverse closest-heavy atom distances were used as input features, of which the importance was normalized. Layer-wise relevance propagation (LRP)^43^ was then applied to the trained network to rank the importance of every input feature for classification. With these approaches, we obtained the importance of every protein residue for distinguishing Nb80-bound and unbound conformational ensembles, and for distinguishing ensembles bound to different ligands. In that way, we could identify which residues were most affected by the binding of the nanobody, and by the binding of different ligands, respectively. As a control, we also calculated the Kullback-Leibler (KL)^23,44^ divergence to derive the important residues. In the KL divergence calculation, high divergences represent non-overlapping inverse distance distributions in active-like states, highlighting thereby features important to distinguish the conformational ensembles.

### Dynamic Network Analysis

Dynamic network analysis for the β2AR in the absence and presence of Nb80 was performed using the *NetworkView* plugin in VMD.^45,46^ For each system, a network map with each protein residue defined as a node was generated. Edges were added between pairs of “in-contact” nodes whose heavy atoms interacted within 4.5 Å for more than 75% of the simulation time. Each edge is weighted by the correction values of two end nodes using the equation: *w_ij_* = *-log*(|*C_ij_*|), in which *w_ij_* and *C_ij_* are the weight and correlation values, respectively. The weight of an edge represents the potential for information transfer (betweenness) between two nodes, where a stronger cross-correlation results in a higher weight, then represented as a thicker edge. Each network was divided into communities of nodes with highly frequent and strong connection to each other using Girvan-Newman algorithm.^47^ Critical nodes that connect the neighboring communities were then identified. The optimal pathways describing the communication between Nb80 and the ligand-binding pocket were also identified based on edges betweenness derived from the correlation of nodes. The Floyd-Warshal algorithm^48^ was used to determine the optimal path between two given nodes: a source and a sink. In general, after the source and sink are chosen, the optimal path is defined to be the connecting route between the two nodes (residues), which minimizes the number of intermediate nodes and maximizes the sum of edge betweenness of the connecting route. In addition, by using the toolkit *subopt*, we identified suboptimal paths, *i.e.* paths that are slightly longer than the optimal path.

## Results and Discussion

### Global structural features derived from data-driven analysis

We used an adaptive sampling protocol to quantitatively sample the most stabilized active-like states of unliganded and ligand-bound β2AR in the absence and presence of Nb80 (**Figure 1 and Table 1**). These conformations are kinetically accessible from the initial active structure (PDB ID 3P0G^25^) and represent snapshots of the protein complex in the active or pre-active states. The ligands studied include the full agonists BI-167107 and adrenaline, the G-protein biased partial agonist salmeterol, the antagonists alprenolol and timolol, and the inverse agonist carazolol. To better understand the receptor conformational changes triggered by binding of Nb80 and various ligands, we performed two different dimensionality reduction analyses: principal component analysis (PCA) and multidimensional scaling (MDS). This allowed us to project the results onto a low dimensional space and clearly visualize overlap between the conformational ensembles. Each point represents a simulation snapshot, which is coloured and marked according to the bound ligand and whether Nb80 is present, respectively (**Figure 2**).

**Figure 2.**
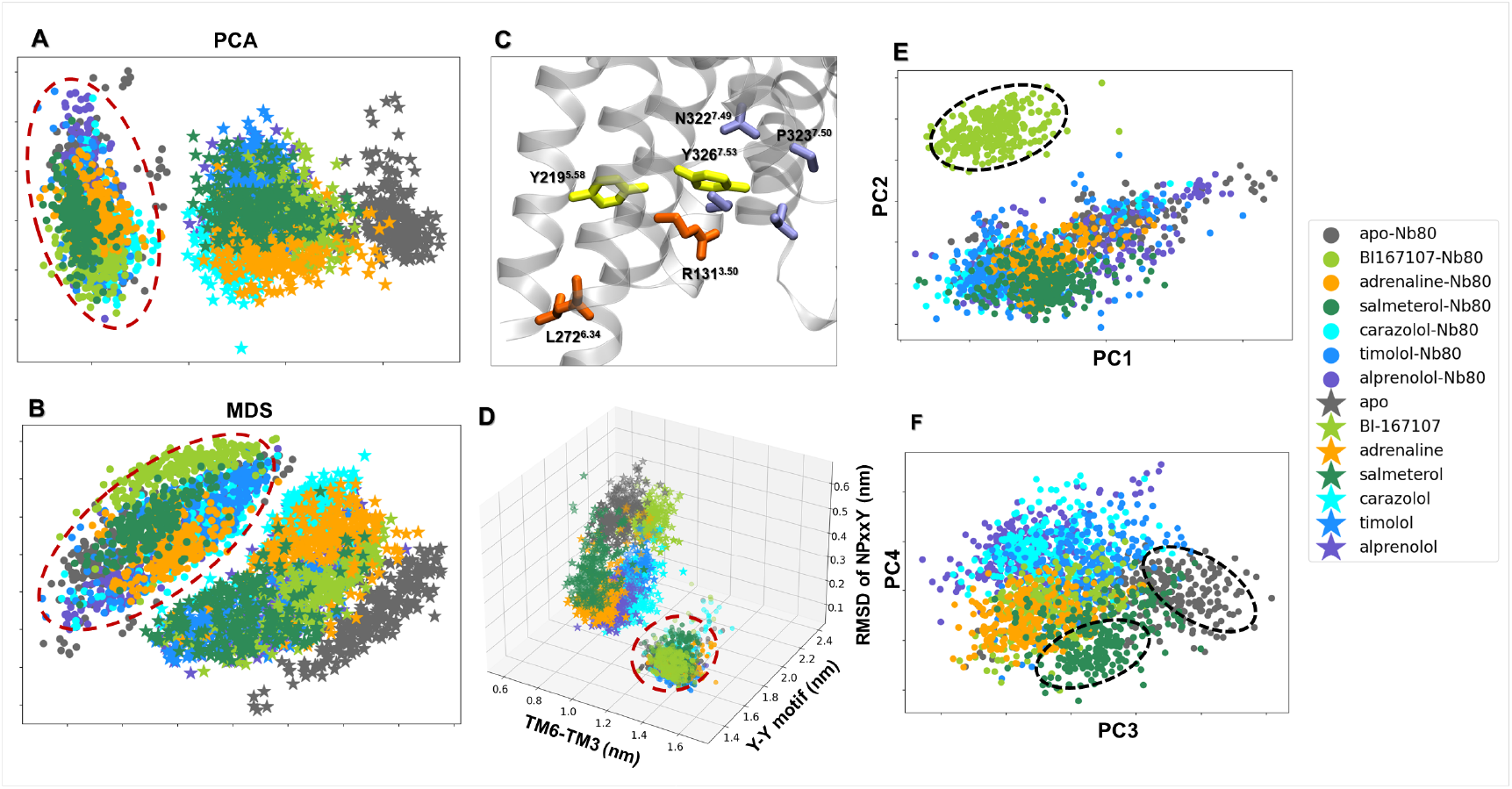
Dimensionality reduction analysis applied to the active-like simulation ensembles. Each point represents a snapshot and is depicted according to the ligand and Nb80-bound ensembles. Input features are the residue-reside Cα atom distances. The Nb80-bound ensembles are highlighted by a red dashed circle. (A) Principal component analysis (PCA) and (B) Multidimensional scaling (MDS) projection on the first two principal components extracted from all trajectories with and without Nb80. (C) Conserved microswitches of the β2AR: R131 and L272 (orange) are located in the transmembrane 3 (TM3) and TM6, respectively. The outward displacement of TM6 is represented by the distance between the Cα atoms of R131^3.50^ and L272^6.34^. Y219^5.58^ and Y326^7.53^ (yellow) are part of TM5 and TM7 which are close to each other via a water-mediated interaction (Y-Y motif) in the β2AR active state. The N^7.59^P^7.50^xxY^7.53^ motif (blue) is on the bottom of TM7. (D) The distributions of the distances between TM6 and TM3, and Y-Y and RMSD of NPxxY motifs. (E-F) PCA projection onto the first four principal components (PC) of Nb80-bound trajectories. BI-167107-, salmeterol-bound and apo snapshots are highlighted by black dashed lines.

#### Nb80-stabilized changes in the β2AR

The PCA and MDS analyses revealed that the Nb80-bound and unbound states are grouped into two distinct clusters (**Figures 2A and 2B**). The absence of overlap between the two clusters for both dimensional reduction methods suggests that Nb80 binding indeed induces conformational changes of the receptor that are independent of ligand binding. In addition, MD snapshots with Nb80 bound tended to be grouped together more compactly than those without Nb80, possibly implying a higher structural rigidity. To intuitively illustrate the Nb80-induced conformational alterations, we further analyzed some traditional microswitches, and measured the outward displacement of transmembrane helix 6 (TM6), the twist of the N^7.49^ P^7.50^ xxY^7.53^ motif and a water-mediated interaction between Y219^5.58^ and Y326^7.53^ (Y-Y motif), which all represent hallmarks of β2AR activation. Here, the TM6 displacement was measured by the distance of Cα atoms between R131^3.50^ and L272^6.34^ (TM6-TM3), and Y-Y motif was represented by the closest heavy atom distance between Y219^5.58^ and Y326^7.53^ (Figure 2C). As shown in Figure 2D, the Nb80-bound ensembles grouped together, away from the Nb80-free states in the 3D distribution space of the distances of TM6-TM3, Y-Y motif and RMSD of NPxxY motif, in agreement with the dimensionality reduction results. Furthermore, for the active-like ensembles without Nb80, binding of different ligands resulted in distinct structural spaces. In contrast, all the snapshots with Nb80 occupied a similar region of the conformational space and gathered into a single cluster represented by an increase in the TM6-TM3 distance, a decrease in the distance of the Y-Y motif, and a decrease in the RMSD of the NPxxY motif. The changes in microswitches indicate a fully active receptor state, especially the outward displacement of TM6, which reaches ~14 Å, similar to the distance measured in the active state structure. This is consistent with a Nb80-mediated enhancement of receptor activation, also reported in previous work.^20^ Our analysis suggests that Nb80 binding triggers conformational changes in the receptor and stabilizes more active-like states, independent of the binding of various ligands. However, no overlap in the projected conformational distribution does not mean that Nb80-bound snapshots are completely different from those without Nb80. There could indeed be overlap in a different projection space. Therefore, we further investigated the conformational space of the third and fourth components for all the ensembles. The result indicates that all simulation snapshots still share common features, whether the Nb80 bound or not (**Figures S1A and S1B**).

#### Ligand-dependent stabilization of β2AR-Nb80 states

The first two-dimensional projections from PCA and MDS as well as the distribution of microswitches highlight the Nb80-induced effects on the receptor conformation, while ligand-mediated structural changes are not resolved in this subspace. To capture the conformational differences between the receptor bound to various ligands, we carried out the same dimensionality reduction methods on the Nb80-bound conformational ensembles only, and projected the conformational ensemble on the first four components, resulting in a different separation of the data (**Figures 2E, 2F, S1C and S1D**). In Figure 2E, BI-167107, a full agonist with ultrahigh affinity to β2AR, segregates away from other ligands, which cluster together. This implies that the binding of BI-167107 induces specific conformational changes in the Nb80-bound receptor. At the same time, the third and fourth components in the projection show many BI-167107 bound snapshots sharing a similar conformational distribution with the others (**Figure 2F**). In addition, we find that most of the unliganded and salmeterol-bound snapshots deviate from the group at the center, indicating that different states are assumed for the two systems (**Figure 2F**).

These results, together with the microswitch conformational distribution, suggest that Nb80 binding promoted all simulation ensembles to share overall features of a fully active state, but the unliganded, BI-167107 and salmeterol stabilized unique activation features. This is in agreement with previous experimental results, supporting the notion that small ligand-specific conformational changes contribute to different receptor activation and downstream signals.^49,50^

### Nb80 and ligand-induced local structural changes

Unsupervised data-driven analysis can provide insights into overall conformational differences of the receptor bound different ligands in the absence and presence of Nb80, but fails to reveal specific Nb80- and ligands-induced activation signatures. To capture important features of receptor activation among active-like states controlled by Nb80 and ligands, we decided to resort to supervised learning methods. We trained classifiers to learn differences between simulation trajectory datasets, using as input inverse inter-residue Ca distances. With this approach, we derive residues that are important to distinguish different receptor-ligand-Nb80 systems, with the idea that these residues could play a substantial role in the receptor activation.

#### Nb80-specific local conformational changes

Importance profiles were calculated using layer-wise relevance propagation on a multilayer perceptron (MLP) classifier trained to distinguish Nb80-bound and unbound states (**Figure 3A**), as well as states modeled in the presence of different ligands from one another, in the presence and in the absence of Nb80 (**Figures 3B and 3C**). As a control, we also characterized important features by computing the Kullback-Leibler (KL) divergence, where residues with high KL divergences are defined as important features (**Figure S2**). Compared to KL divergence, the MLP classifier generates importance profiles with more peaks, as it can find all important features by performing nonlinear transformations of input features.^23^ We observed that both methods identified the cytoplasmic end of TM6 as the most important region to discriminate states with and without Nb80 (**Figures 3A and S2A**). Recent studies have identified multiple inactive, intermediate and active receptor states with different degrees of conformational changes at the intracellular end of TM6, in which complete receptor activation accompanied by a ~14 Å outward movement of TM6 requires both agonist and G protein or a mimetic nanobody such as Nb80.^20,51^ Notably, this region does not differentiate the ligand-bound receptor modeled in the presence or absence of Nb80 (**Figures 3B and 3C**), suggesting that the cytoplasmic end of TM6 conformations are very similar within these two classes. This is also compatible with the conformational distribution of microswitches (**Figure 2D**). In addition, the MLP classifier also highlighted some residues on TM3, TM5 and TM7, which exhibited different conformations in the Nb80-bound and -unbound states.

**Figure 3.**
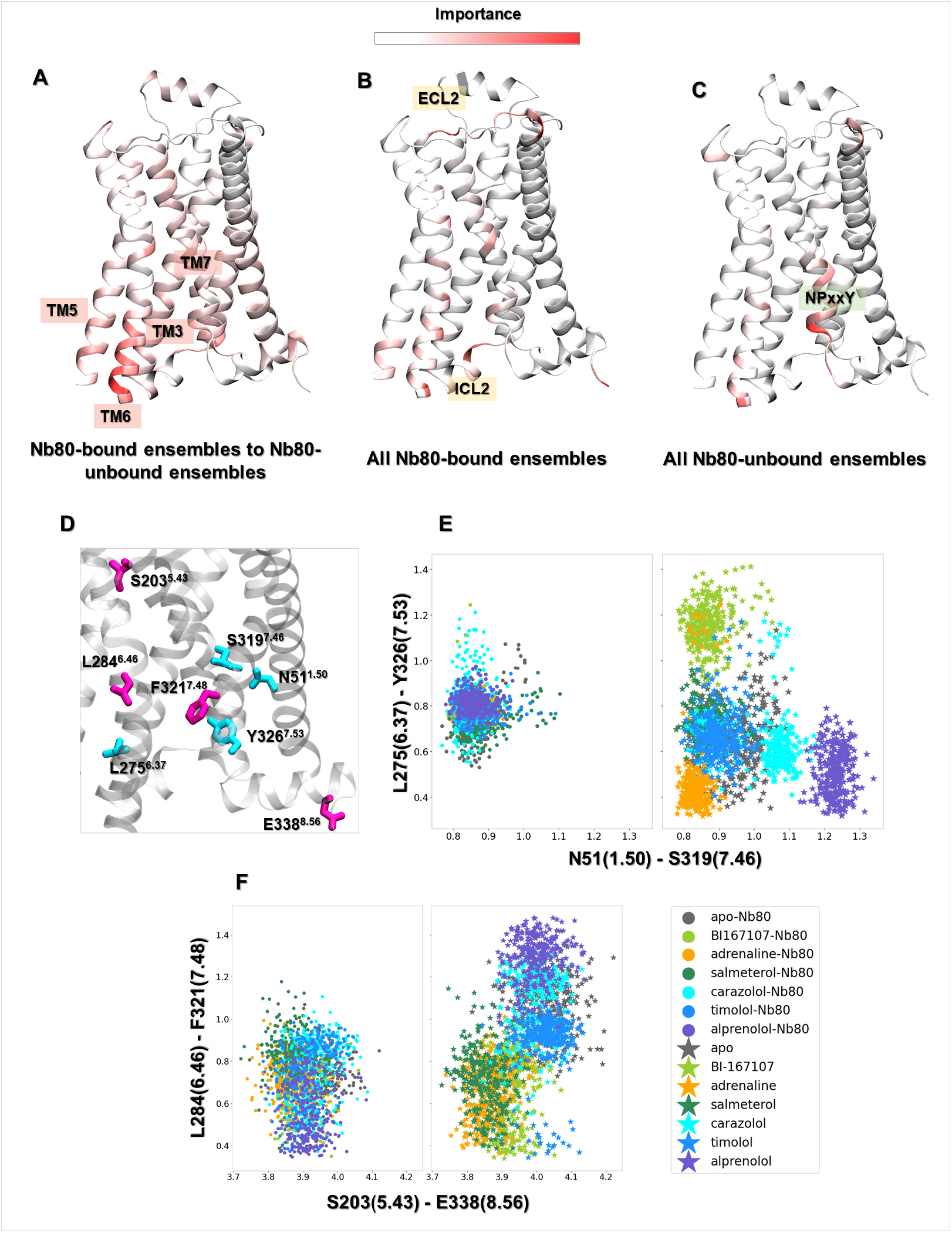
Residues important for discriminating Nb80-dependent activation mechanisms, derived from training a multilayer perceptron (MLP) classifier on equilibrated active-like ensembles. (A) Comparison between Nb80-bound and Nb80-unbound ensembles. The most important hotspot is located at the end of TM6. (B) Residues important to distinguish the Nb80-bound ensembles. (C) Residues important to distinguish the Nb80-unbound ensembles. (D) Residues important to differentiate Nb80-bound from Nb80-unbound ensembles. (E) Distribution of N51^1.50^-S319^7.46^ and L275^6.37^-Y326^7.53^ distances. (F) Distribution of S203^5.43^-E338^8.56^ and L284^6.46^-F321^7.48^ distances.

In contrast, there are only a few identified hotspots for discriminating all β2ΑR-Nb80 complexes, illustrating that all the bound ligands stabilized common structural rearrangements (**Figure 3B**). Among the few regions distinguishing ligand-bound ensembles, a few residues in the intracellular loop (ICL) 2 and extracellular loop (ECL) 2 showed up as important features when comparing receptor ensembles when bound to the various ligands in the presence of Nb80. In agreement with this, several studies point out that ECL2 is involved in ligand specificity, and in determining the affinity of ligands towards the receptor.^52^ Moreover, it should be stressed that ICL2 directly interacts with the N-terminus of G-protein and is responsible for the selectivity of receptor - G-protein interactions as well as the efficiency of G-protein activation.^53–55^ Besides, for all receptor states without Nb80 bound, the NPxxY motif exhibited a ligand-specific conformation, in agreement with our previous study^27^ (**Figure 3C**). In contrast, the NPxxY motif adopts a similar conformation for all ligand-bound ensembles in the presence of Nb80 (**Figures 3B and 2D**).

In addition to conformational differences in the cytoplasmic region induced by Nb80 binding, structural changes through the TM domain were also captured by this analysis. Several key residues with a higher importance for distinguishing Nb80-bound active-like states were extracted for further investigation (**Figure 3D**). For the N80-free states, agonists governed different TM6 and TM7 orientations near the NPxxY motif, leading to distinct distances between Y326^7.53^ and L275^6.37^. In the same region, a hydrogen bond formed between S319^7.46^ and N51^1.50^, one of the most conserved residues in the class A GPCRs, only in the agonist-bound receptor. However, we notice that Nb80 binding stabilized similar conformation between Y326^7.53^ and L275^6.37^ and maintained the hydrogen contact of S319^7.46^ with N51^1.50^ regardless of ligand bound (**Figure 3E**). Moreover, our analysis indicated that agonists induced a local contraction between L284^6.46^ and F321^7.48^ and a long-range contraction between S203^5.43^ and E338^8.56^ compared to non-agonists in the absence of Nb80. These residues are located around TM5 bulge, PIF motif, and NPxxY motif, and play an important role in the receptor activation (**Figure 3D**). However, from the data-driven analysis, the binding of Nb80 could make the distribution for the above four residues overlap for non-agonists-bound receptor features (**Figure 3F**). Such comparison further supports the finding that Nb80 binding induces some structural rearrangements throughout the protein and stabilizes an active conformation of the β2AR independently of the chemical nature of the ligand bound in the extracellular site. This also suggests a higher free energy barrier for Nb binding for non-agonist ligands.

#### Ligand-specific local conformational changes in the presence of Nb80

To better understand the different ligand-induced conformational changes in β2ΑR-Nb80 complexes, the receptor ensembles of the apo, BI167107-bound, and salmeterol-bound, which all occupied a distinct region of the conformational space in the dimensional reduction analysis (**Figures 2E and 2F**), were labeled as separate datasets for further classification. Interestingly, compared with the full agonist BI167107, salmeterol is a functionally selective partial agonist for β2ΑR with a 5- to 20-fold bias toward G-Protein Gs over arresti.^25,56^ Also, its high selectivity and long-acting properties contribute to it being one of the most prescribed drugs for treating asthma and chronic obstructive pulmonary disease (COPD).^57^ Using a similar protocol as above, we identified features specific to the three chosen ensembles (**Figure 4**). Compared to the two ligand-bound ensembles, there are more important residues located in the TM domain in the unliganded ensemble (**Figures 4A, 4B and 4C**). This may originate from the large flexibility of the apo receptor (**Figure S3**), in agreement with spectroscopy experiments suggesting that the β2ΑR could not be stabilized in its fully active state in absence of agonist binding.^20^

**Figure 4.**
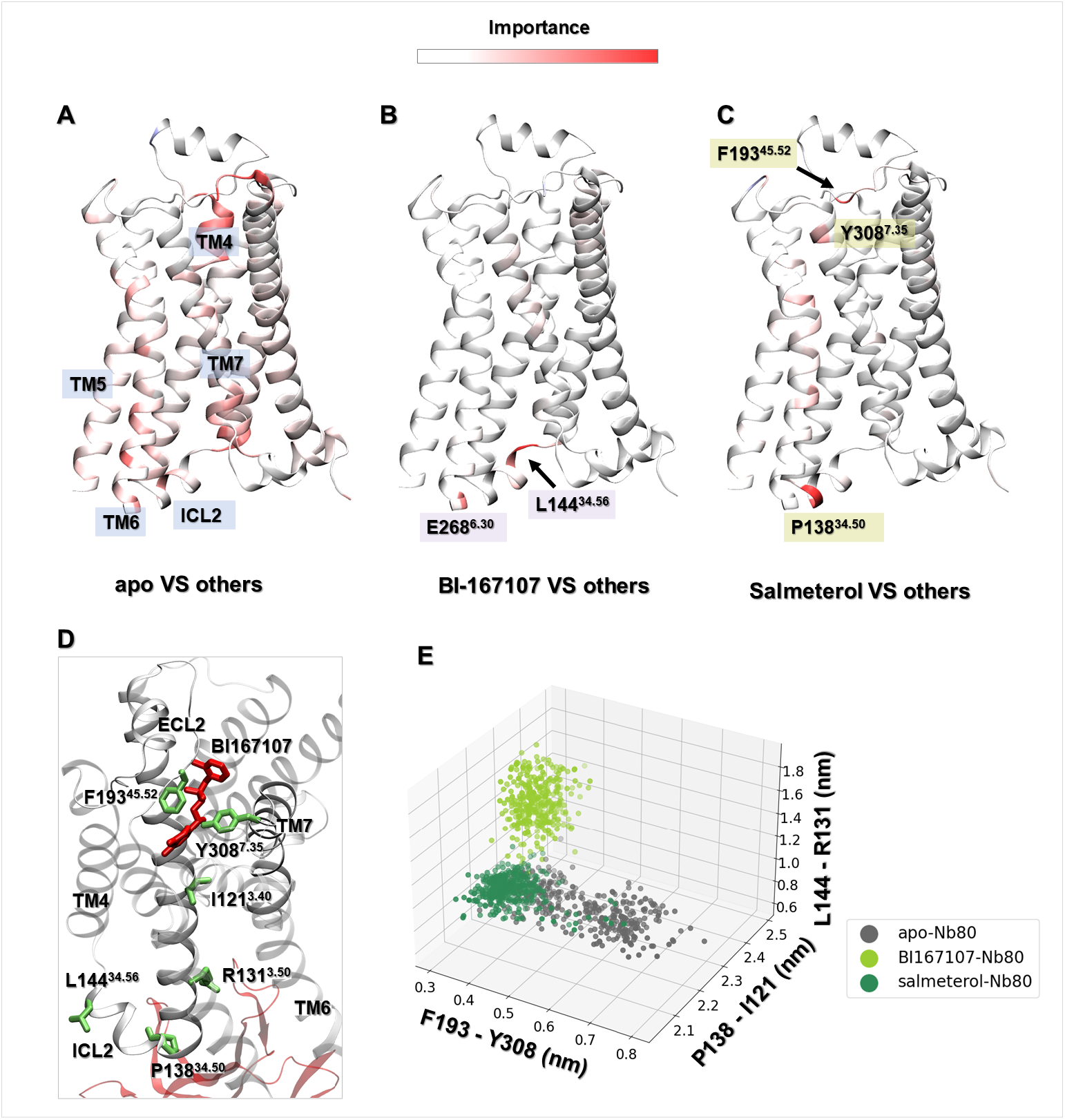
Important residues derived from the equilibrated active-like ensembles for discriminating ligand-dependent activation mechanisms using a multilayer perceptron (MLP) classifier. (A-C) Comparison of the apo, BI167107-, and salmeterol-bound ensembles to the others respectively in the presence of Nb80. (D) Important residues for differentiating apo, BI167107- and salmeterol-bound β2ΑR-Nb80 ensembles. (E) Distances distribution between F193^45.52^ - Y308^7.35^, P138^34.50^ - I121^3.40^ and L144^34.56^ - R131^3.50^ in apo, BI-167107- and salmeterol-bound β2ΑR-Nb80 ensembles.

Notably, only a few residues in the BI-167107- and salmeterol-β2AR-Nb80 complexes were captured as important, which indicates that there were only subtle differences between their conformational ensembles. Those corresponded to, for example, L144^34.56^ and E268^6.30^ in the BI-167107-bound state, and P138^34.50^, F193^45.52^ and Y308^7.35^ in the salmeterol-bound state. Among them, L144^34.56^ and P138^34.50^ are located in ICL2, which is associated with distinct ligand-dependent conformational changes to recognize G-proteins or β-arrestin.^54,58^ Furthermore, mutational and biophysical analysis suggested that F193^45.52^ and Y308^7.35^ are closer to each other in the agonist-bound β2AR-Nb80 complex and form a lid-like structure over the orthosteric ligand-binding pocket, which slowed down the rate of ligand dissociation, and accordingly contributed to the enhancement of the ligand affinity.^49^ As shown in Figure 4E, the distance between F193^45.52^ and Y308^7.35^ in unliganded simulation snapshots ranged from 3 Å to 7 Å, while in BI167107- and salmeterol-bound states it was stabilized around 3-4 Å, which provides a structural explanation for the agonist-induced enhancement of receptor activation observed in experiments.^59,60^ Meanwhile, we noticed that the F193^45.52^ - Y308^7.35^ distance in salmeterol-β2AR-Nb80 complex is slightly larger than that in BI167107-β2AR-Nb80 complex, which can be related to the lower affinity and partial activation effect of salmeterol. Moreover, mutagenesis studies^33,61^ have reported that the hydrogen bond between F193^45.52^ and the aryl-oxy-alkyl tail of salmeterol contributed to its high selectivity for β2ΑR over β1ΑR.

Furthermore, we also identified other residue-pairs showing different conformations induced by BI167107 and salmeterol binding (**Figures 4D and 4E**). Among them, I121^3.40^ and R131^3.50^ are part of the PIF motif and the “ionic lock”, respectively, which are hallmarks of GPCR activation.^62^ We observe shorter distances of the residues pairs P138^34.50^ and I121^3.40^, and L144^34.56^ and R131^3.50^ in the salmeterol complex than in the BI167107 complex, indicating a loose interaction connecting the intracellular region and ligand-binding site in the BI167107-bound structure. Overall, there are indeed some distinct structural features associated with the receptor activation presenting in Nb80-stabilized β2ΑR bound to ligands with varying efficacies.

In general, our approach has succeeded in identifying important features distinguishing Nb80-bound and -unbound states. In addition to the intracellular end of TM6, some highly conserved residues such as N51^1.50^, S319^7.46^, S203^5.43^ and Y326^7.53^ were identified to play crucial roles in the receptor activation. ICL2, involved in G-protein activation, was also highlighted to be important in different ligands-bound β2ΑR-Nb80 structures. Interestingly, F193^45.52^ was captured as a key factor for the selectivity of salmeterol in β2ΑR activation.

### Dynamic allosteric network in the β2ΑR

To gain insights into the allosteric communication pathways modulated by Nb80 and ligands, we constructed a residue interaction network model using the MD simulation ensembles of the β2ΑR and analyzed it using community network analysis (see Methods). We specifically focused on six ensembles, including unliganded, BI167107- and salmeterol-bound β2ΑR in the presence and absence of Nb80 (**Figures 5 and S4**). As shown in **Figure 5**, there are distinct intercommunity flows in these different receptor states. Overall, a smaller number of communities are identified in the Nb80-bound structures compared to those in Nb80-free states, indicating that Nb80 binding induces tighter and stronger local communication networks and consequently bigger communities. For example, in the unliganded structures, Nb80 binding promoted the grouping of communities 5, 9 and 10 (C5, C9 and C10), which are mainly found at the intracellular end of TM3, TM5, TM6 and ICL2, which correspond to a bigger community C5 in the apo-β2ΑR-Nb80 complex (**Figures 5A and 5D**). Similarly, communities C1 and C7 located at the extracellular domain of the BI167107-bound structure merged into a single cluster C1 upon binding of Nb80 (**Figures 5B and 5E**). However, the dynamic network of the salmeterol-bound receptor is different from the others in the presence of Nb80, especially in the extracellular region (**Figure 5C**). We observed that community C1 in the unliganded and BI167107-bound states was split into C1 and C9 in the salmeterol-bound state. This might be attributed to the long aryl-oxy-alkyl tail of salmeterol, which led to the generation of an exosite consisting of residues around ECL2, ECL3, and the extracellular ends of TM6 and TM7. Interestingly, the exosite is associated with high receptor selectivity and ligand affinity.^33^

**Figure 5.**
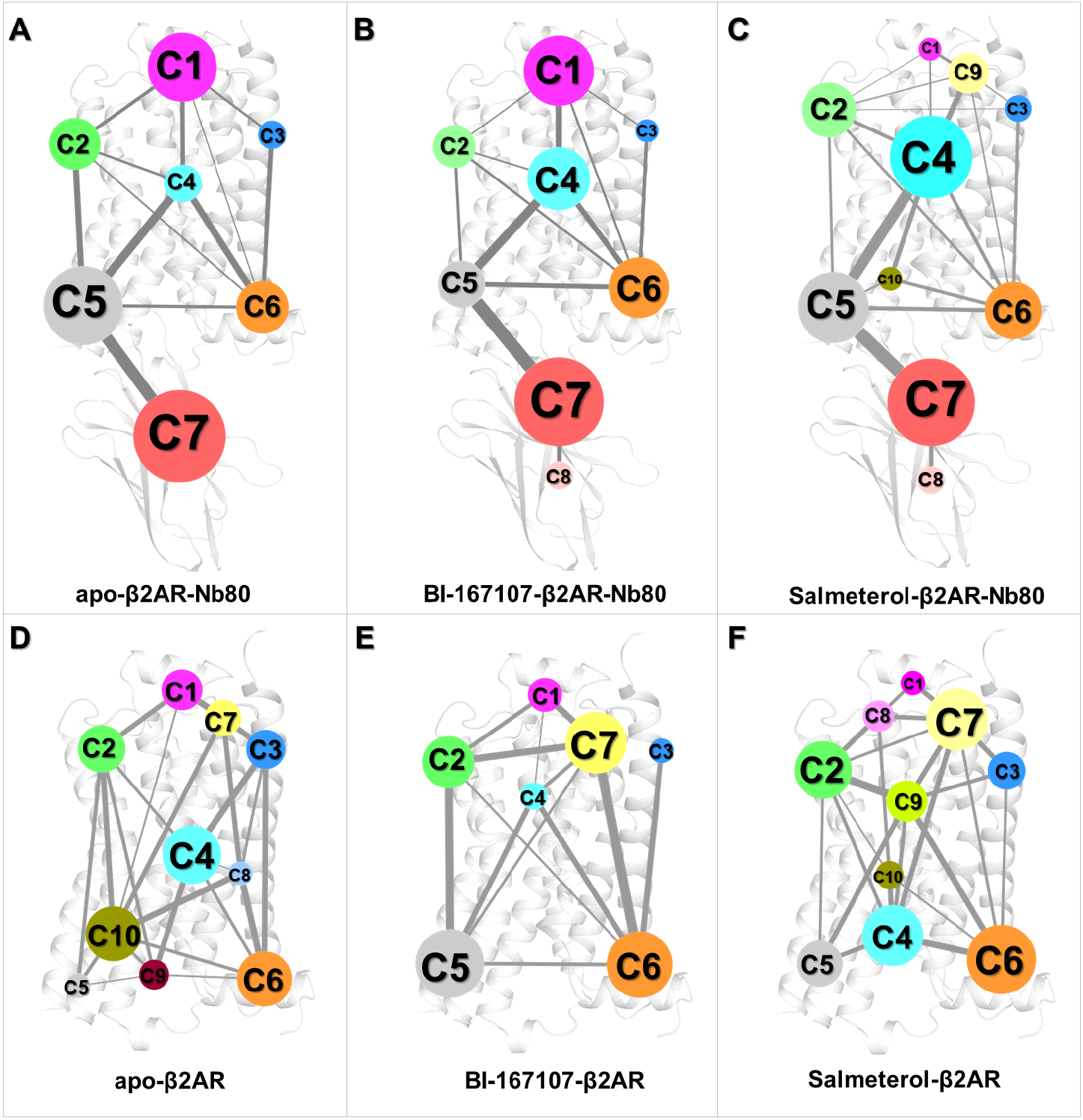
Dynamic networks of the apo, BI167107- and salmeterol-bound β2ΑR with and without Nb80 bound are analyzed using community network analysis. (A-C) 2D networks of unliganded, BI167107- and salmeterol-bound forms with Nb80. (D-F) 2D networks of apo, BI167107- and salmeterol-bound forms without Nb80. Network communities are colored separately according to their ID number. A community represents a set of highly intra-connected nodes (residues), its size being determined by the number of nodes included in a community. Edges connecting two communities are represented by lines, of which the width is proportional to the strength of the information flow between the connected communities.

In addition, nodes (residues) critical for the communication across communities were identified for the three receptor-Nb80 ensembles (**Figure 6**). Residues R131^3.50^, I127^3.46^, I112^3.31^, L115^3.34^ located on TM3, which is an important signal transduction domain across class A GPCRs,^63^ were identified as critical residues in all the Nb80-bound structures. Furthermore, other critical residues like D192^45.51^ and K305^7.32^ in BI167107- and salmeterol-bound states have been reported to contribute to the formation of a closed conformation over the ligand-binding pocket, in part responsible for enhanced ligands binding affinity.^49^ We also found some ligand-specific critical residues, such as K97^2.68^, E306^7.33^, F290^6.52^, and Y209^5.48^ in the BI167107-bound state and C184^45.43^, C191^45.50^, W313^7.40^ and S207^5.46^ in the salmeterol-bound state, which also have an effect on the receptor activity and ligand affinity, supported by the mutagenesis data reported in the G-protein-coupled receptor data bank (GPCRdb, http://gpcrdb.org).^64,65^

**Figure 6.**
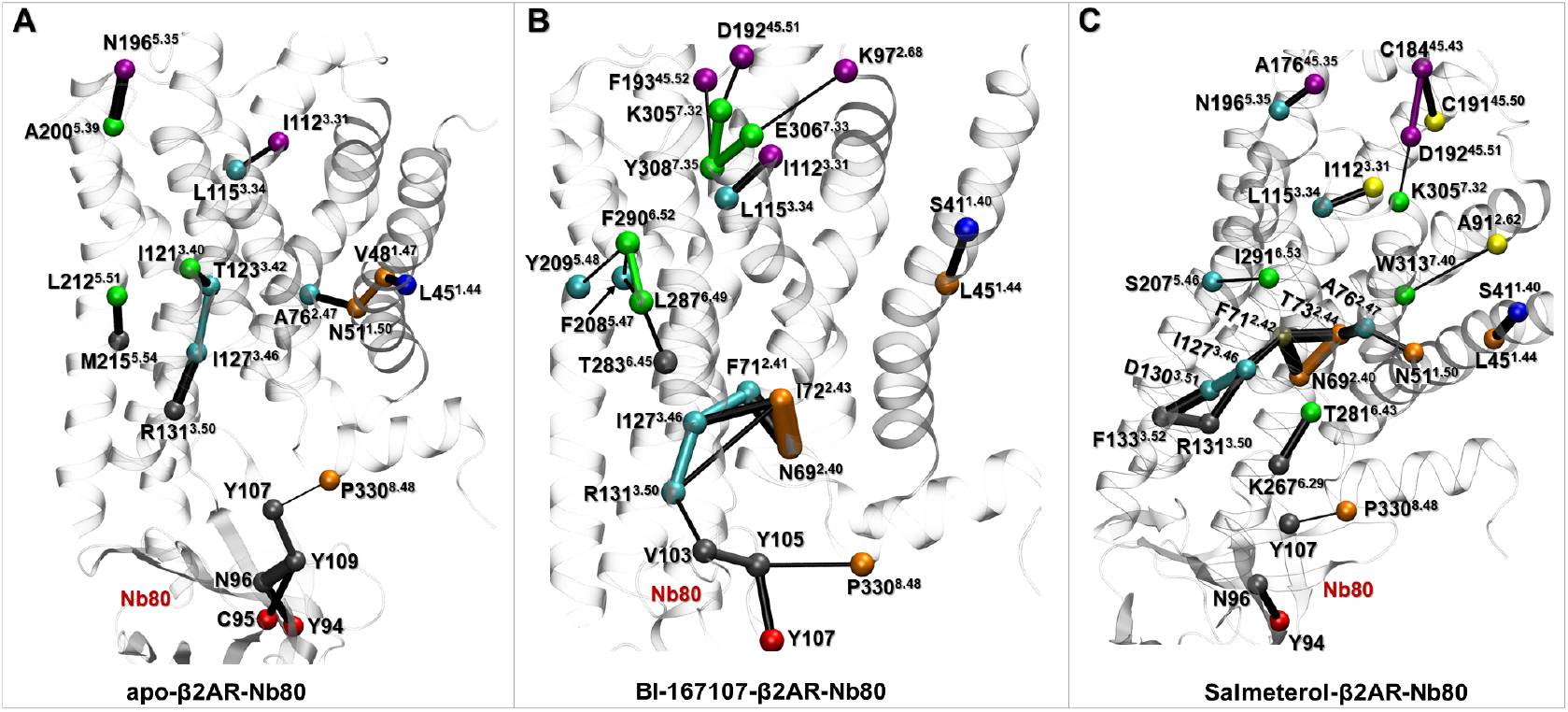
Critical nodes in the apo, BI-167107- and salmeterol-bound β2ΑR-Nb80 structures. Each critical node is located at the interface of neighboring communities, and corresponds to the edge with the highest score in terms of connectivity. Critical nodes are colored consistently with the communities of dynamic network models of Figure 5, and the connecting edges are represented by lines with their width weighted by betweenness.

We further analyzed the optimal pathways in the three Nb80-bound structures, to identify residues involved in information transfer from the Nb80-binding site to the extracellular domain of the receptor (**Figure 7**). For each network model, we selected critical nodes in communities C7 (Nb80) and C1 (extracellular binding site) as start- and end-points, respectively, for pathways calculation. Those were Y94, N196^5.35^, L112^3.31^ and H93^2.64^ in the unliganded state, Y107, F193^45.53^, C192^45.51^ and K97^2.68^ in the BI167107-bound state, and Y94, A176^45.35^, C184^45.44^ and C192^45.51^ in the salmeterol-bound state (**Figure 6**). The C5 community contains residues from both the Nb80 and intracellular ends of TM3, TM5 and TM6, forming an interfacial community. More residues of Nb80 merged into C5 in the unliganded and salmeterol-bound networks than in the BI167107 bound network. The unliganded and salmeterol-β2ΑR-Nb80 states share the inner Nb residue Y94 as an important residue for communication between the Nb80-only community C7 with the mixed receptor-Nb80 community C5. In contrast, in the BI167107-bound ensemble, the surface residue Y107 fulfills this role. This suggests a comparatively loose interaction induced by BI167107 at the β2ΑR-Nb80 interface (**Figure 7 and S5**).

**Figure 7.**
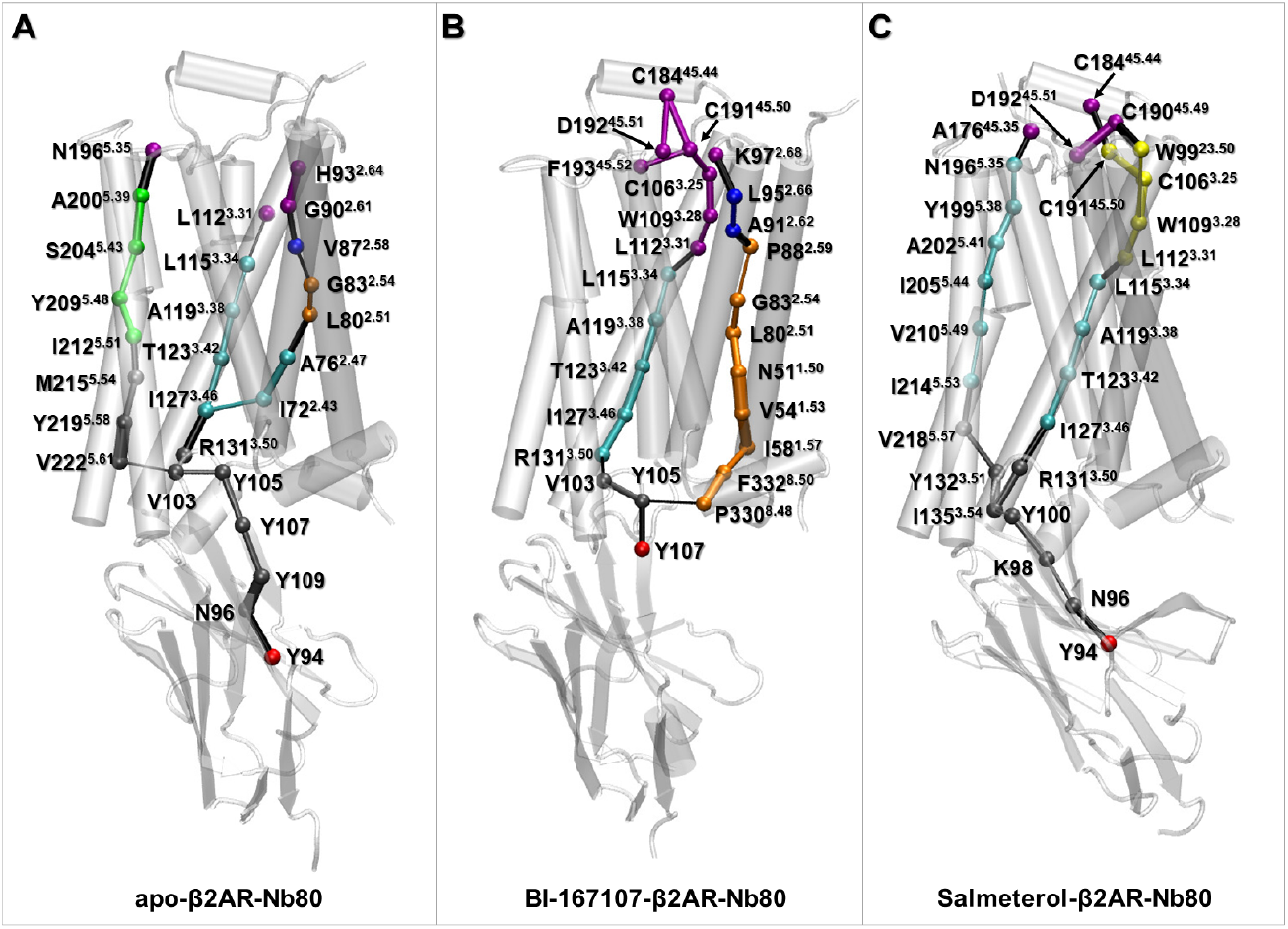
Optimal paths connecting the intracellular (C7) and extracellular binding sites (C1) in apo (A), BI167107 (B), and salmeterol (C) -bound β2ΑR-Nb80 structures. Residues are rendered as spheres and colored consistently with the communities they belong to in Figure 5, and the connecting edges are represented by lines with their width weighted by betweenness.

In addition, the major difference in the three models is that there are three optimal pathways connecting the extra- and intracellular binding sites in the unliganded structure, while there are only two in ligand-bound structures. This could be expected from the more prominent fluctuations in the ligand-free receptor (**Figure S3**). Indeed, we observed one pathway going primarily along TM3 in all three network models, which used R131^3.50^ of the ionic lock as a bridge node connecting the Nb80-binding and ligand-binding sites. This is supported by previous studies emphasizing the significant role of TM3 in signal transduction between the intracellular and extracellular binding sites.^63,66^ Notably, the BI167107-specific pathway sent signals mainly along H8, TM1 and TM2, whereas the salmeterol-bound receptor’s pathway prominently involved TM5 (**Figures 7B and 7C**). Several studies pointed out that TM2 might be regarded as a pivot for activating conformational change of GPCRs, in which the Pro residue at 2.58, 2.59 or 2.60 may contribute to specialize GPCRs binding of different ligand types (P88^2.59^ in the β2ΑR).^63,67^ In addition to P88^2.59^, N51^1.50^ in BI167107-specific optimal pathway is associated with water-mediated interactions around the cytoplasmic halves of TM2, TM6 and TM7, playing a crucial role in GPCR activation.^68^ In the case of the salmeterol-specific pathway, I205^5.44^ and V210^5.49^ are located near S207^5.46^ (the TM5 bulge) and I211^5.50^ (PIF motif), which are involved in highly conserved microswitches.^62^ The hydrophobic interactions involving I205^5.44^ and V210^5.49^ may indirectly help stabilize the inward conformations of S207^5.46^ and I211^5.50^. Moreover, V218^5.57^ and N196^5.35^ contributed to signal transmission connecting the intracellular end of TM3 and ECL2 region (**Figure 7C**).

To summarize, network analysis revealed that Nb80 induced high levels of communication especially in the intracellular domains of TM3, TM5, TM6 and ICL2, and extracellular domains of TM2, TM3, TM5, TM7 and ECL2. With this approach, we also identified critical residues that had important effects on the receptor activity and ligand affinity. In addition, ligands-specific allosteric signaling pathways highlighted different conformational changes controlled by the ligands.

## Conclusion

Many studies^49,50,69,70^ have shown that nanobodies, functioning as G-protein mimetics, succeed in stabilizing different GPCR conformations and further affect the affinity of ligands by allosteric modulation. Nb80, the first reported nanobody, bound intracellularly to β2ΑR not only stabilizes the active agonist-bound receptor conformation but also highly improves the agonist affinity. In this work, on the basis of data-driven methods and dynamic network analysis for active-like β2ΑR ensembles bound to ligands with varying efficacies in the absence and presence of Nb80, we propose a molecular interpretation of the allosteric modulation mechanism due to Nb80 binding. Nb80 binding was found to stabilize the same conformational rearrangements for different systems, especially the larger intracellular outward movement of TM6 and the decrease in the distance of the Y-Y motif and the RMSD of NPxxY motif. Highly conserved residues N51^1.50^, S319^7.46^, S203^5.43^, L284^6.46^ and Y326^7.53^ are identified to be important in the Nb80-stabilized active β2ΑR conformation. Network analysis further reveals Nb80-induced stronger interactions in the intracellular and extracellular domains of the receptor. In addition, apo, BI167107- and salmeterol-bound states exhibit subtle differences in TM3, ECL2 and ICL2 induced by resides such as I121^3.40^, R131^3.50^, F1934^5.52^, P1383^4.50^ and L1443^4.56^, some of which are also identified as critical nodes in dynamical network models and proved to be important for the receptor activity by previous mutagenesis experiments.^62–64^ Interestingly, we observed that the BI167107- and salmeterol-specific optimal pathways contribute to the signal transmission connecting Nb80 and ligand binding sites mainly via TM1 and TM2, and TM5 respectively.

Thus, enhanced sampling MD simulations combined with data-driven analysis methods were useful to probe the allosteric effect of Nb80 binding. Our results shed light on ligands-specific subtle structural differences and signal transmission pathways. This work provides structural insights underlying the enhanced β2ΑR activation activity and ligand affinity modulated by Nb80. These findings could be helpful for structure-based drug discovery targeting GPCRs, taking into account the effect of intracellular binding partners.

## Supporting Information

The supporting information is available free of charge on the ACS Publications website at DIO: 10.1021/xxxxx.

Dimensionality reduction analysis applied to the active-like simulation ensembles; Important residues derived from the equilibrated active-like ensembles for discriminating Nb80- and ligand-dependent activation mechanisms by computing Kullback-Leibler divergence (KL); Residue average fluctuations measured as root-mean-square fluctuation (RMSF) in the active-like simulation ensembles of apo, BI167107- and salmeterol-bound β2ΑR-Nb80 structures; Dynamic networks are identified in the apo, BI167107- and salmeterol-bound β2ΑR with and without Nb80 bound through community network analysis; Local network communities involve Nb80 and the intracellular domain of β2AR.

The data necessary to reproduce the findings presented in this paper can be found on OSF (DOI 10.17605/OSF.IO/6XPYV). The code used to run and analyze simulations has been deposited on GitHub (https://github.com/delemottelab/demystifying, and https://github.com/delemottelab/state-sampling)

## Notes

The authors declare no competing financial interest.

