## Supplementary Figures for "Allosteric effect of nanobody binding on ligand-specific active states of the β2-Adrenergic Receptor"

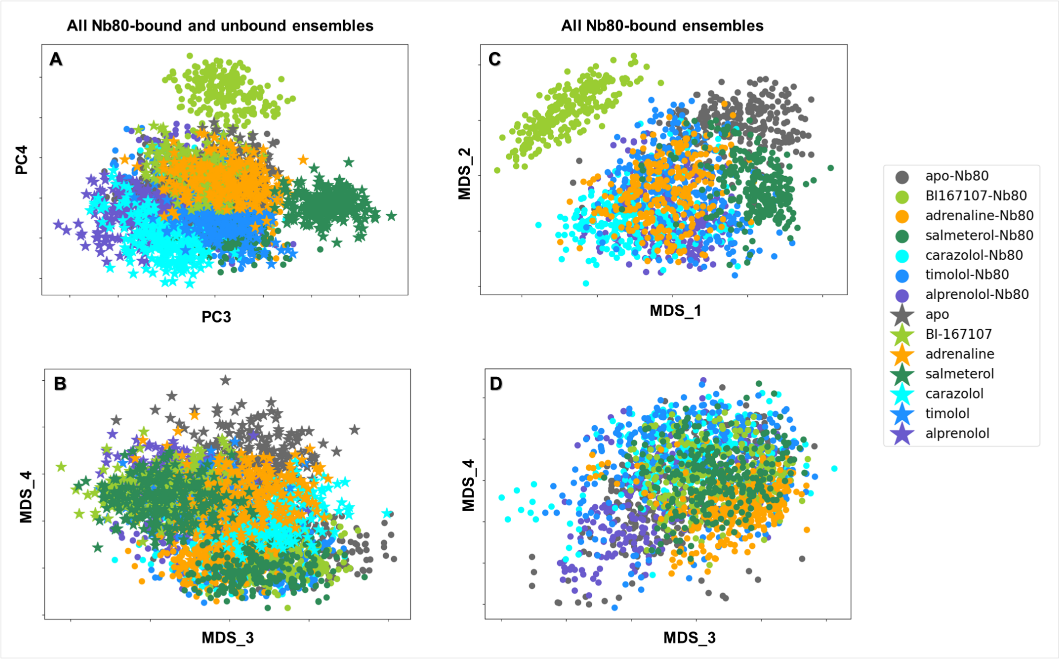


**Figure S1.** Dimensionality reduction analysis applied to the active-like simulation ensembles. Each point represents a simulation snapshot and is colored according to the ligand and Nb80 bound ensembles. (A-B) PCA and MDS projection onto the third and fourth components of all trajectories with and without Nb80. (C-D) MDS projection on the first four components of trajectories with Nb80.

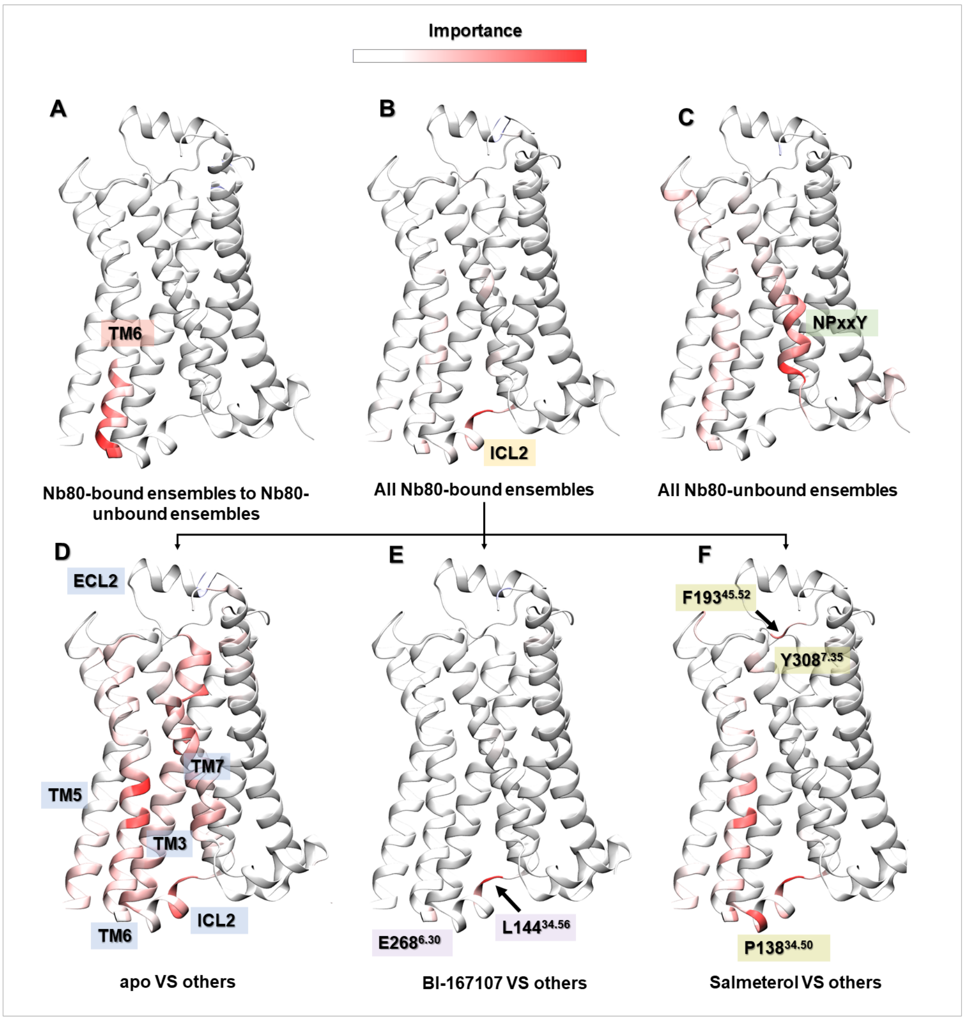


**Figure S2**. Important residues derived from the equilibrated active-like ensembles for discriminating Nb80- and ligand-dependent activation mechanisms by computing Kullback-Leibler divergence (KL).

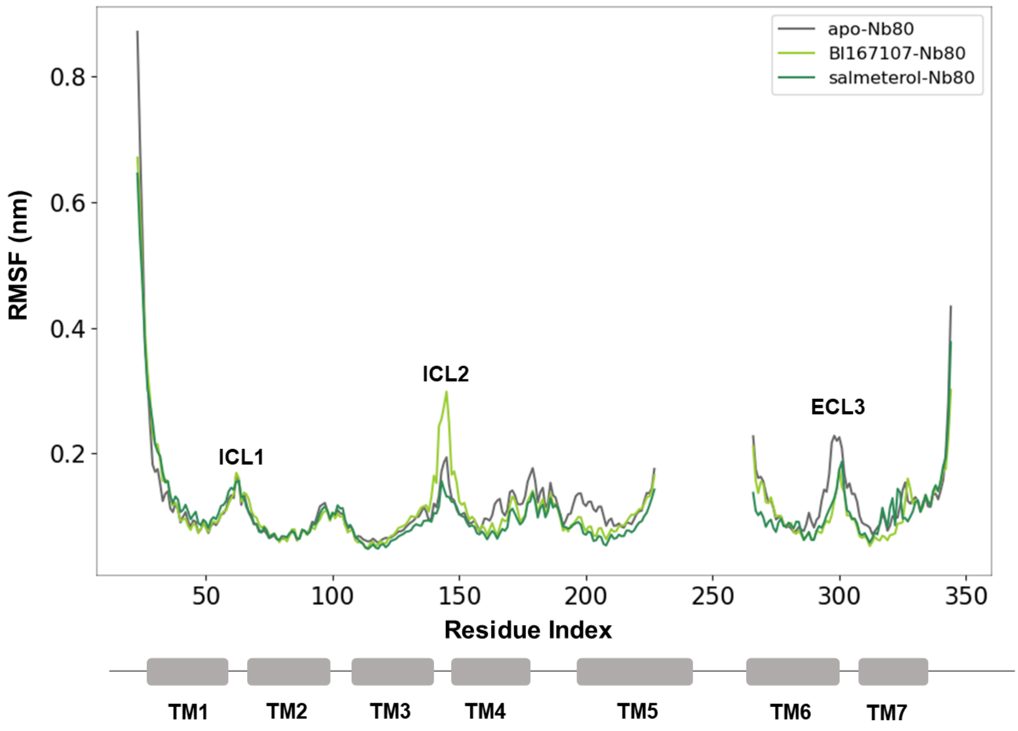


**Figure S3.** Residue average fluctuations measured as root-mean-square fluctuation (RMSF) in the active-like simulation ensembles of apo, BI167107- and salmeterol-bound β2AR-Nb80 structures.

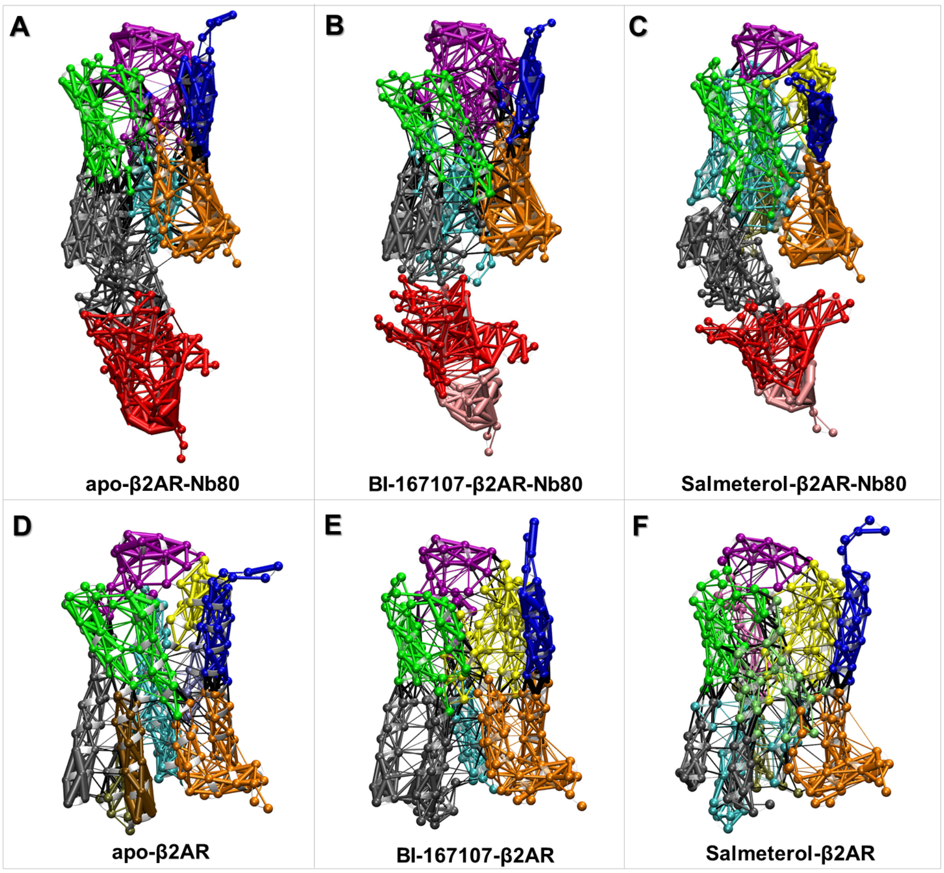


**Figure S4**. Dynamic networks are identified in the apo, BI167107- and salmeterol-bound β2AR with and without Nb80 bound through community network analysis. (A-C) 3D networks of unliganded, BI167107- and salmeterol-bound forms with Nb80. (D-F) 3D networks of unliganded, BI167107- and salmeterol-bound forms without Nb80. Network communities are colored separately by their ID number. Residues are rendered as spheres and the connecting edges are represented by lines with their width weighted by betweenness.

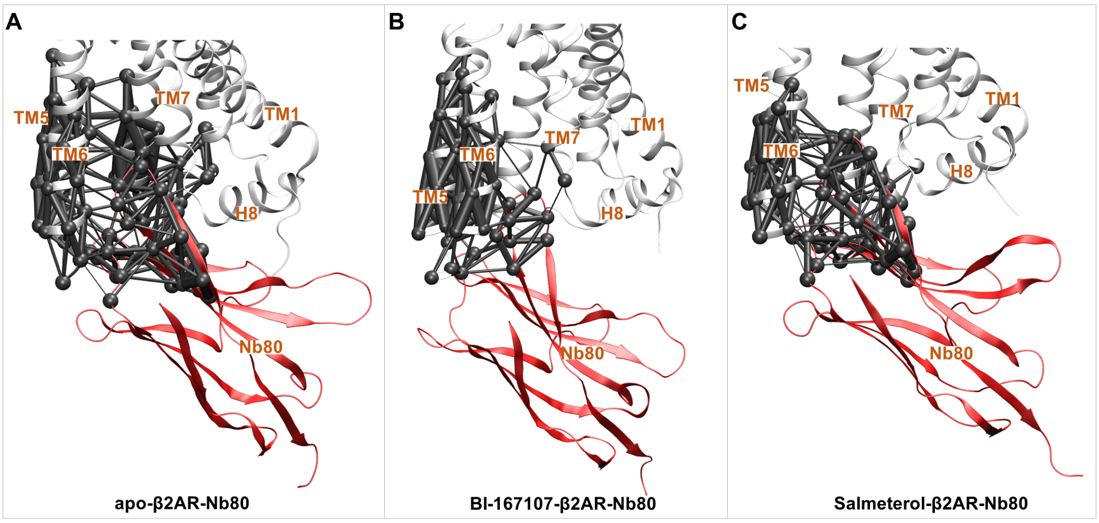


**Figure S5.** Local network communities involve Nb80 and the intracellular domain of β2AR. β2AR and Nb80 are represented by white and red cartoons respectively. Gray community in the BI167107-β2AR-Nb80 system contains less residues of Nb80 than those in apo and salmeterol-β2AR-Nb80 systems. Residues in the network are rendered as spheres, and the connecting edges are represented by lines with their width weighted by betweenness.
